# Spatial determination and prognostic impact of the fibroblast transcriptome in pancreatic ductal adenocarcinoma

**DOI:** 10.1101/2023.01.20.524883

**Authors:** Wayne Croft, Hayden Pearce, Sandra Margielewska-Davies, Lindsay Lim, Samantha M. Nicol, Fouzia Zayou, Daniel Blakeway, Francesca Marcon, Sarah Powell-Brett, Brinder Mahon, Reena Merard, Jianmin Zuo, Gary Middleton, Keith Roberts, Rachel M. Brown, Paul Moss

**Affiliations:** Institute of Immunology and Immunotherapy, College of Medical and Dental Sciences, University of Birmingham, Birmingham, UK; Centre for Computational Biology, University of Birmingham, Birmingham, UK; Cancer Research Horizons, The Francis Crick Institute, London, UK; University Hospitals Birmingham NHS Foundation Trust, Queen Elizabeth Hospital Birmingham, Birmingham, UK

**Keywords:** Pancreatic cancer, fibroblasts, spatial, prognosis

## Abstract

Pancreatic ductal adenocarcinoma has a poor clinical outcome and responses to immunotherapy are suboptimal. Stromal fibroblasts are a dominant but heterogenous population within the tumor microenvironment and therapeutic targeting of stromal subsets may have therapeutic utility. Here we combined spatial transcriptomics and scRNA-Seq to define the transcriptome of tumor-proximal and tumor-distal cancer-associated fibroblasts (CAFs) and linked this to clinical outcome. Tumor-proximal fibroblasts comprised large populations of myofibroblasts, strongly expressed podoplanin, and were enriched for Wnt ligand signaling. In contrast, inflammatory CAFs were dominant within tumor-distal subsets and expressed complement components and the Wnt-inhibitor SFRP2. Poor clinical outcome was correlated with elevated HIF-1α and podoplanin whilst high level expression of inflammatory and complement genes was predictive of extended survival. These findings demonstrate the extreme transcriptional heterogeneity of CAFs and its determination by apposition to tumor. Selective targeting of tumor-proximal subsets, potentially combined with HIF-1α inhibition and immune stimulation, may offer a multimodal therapeutic approach for this disease.

**Graphical Abstract:** 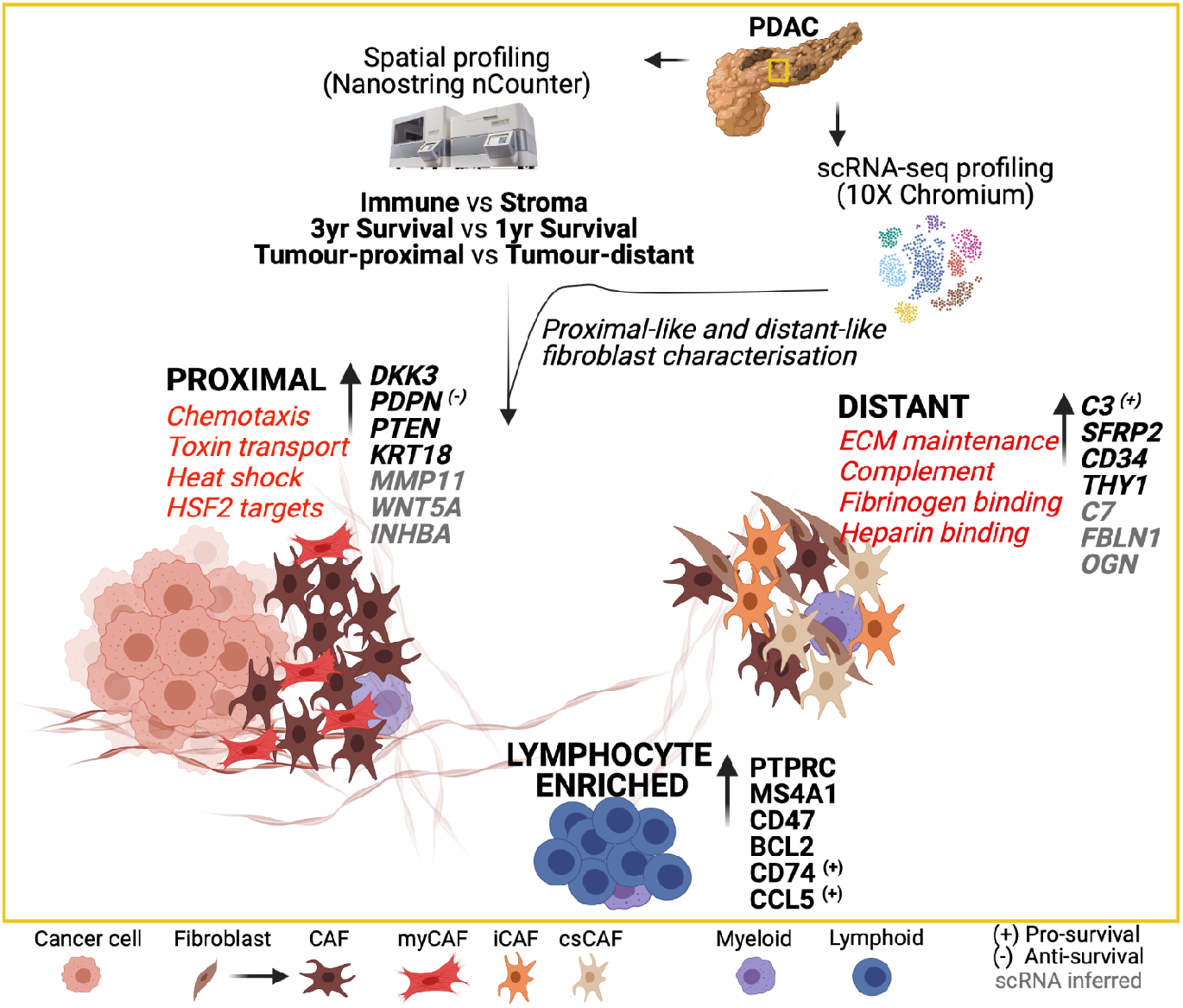

## Introduction

Therapeutic control of pancreatic ductal adenocarcinoma (PDAC) is one of the greatest challenges in oncology and PDAC remains associated with poor long term survival (1). Although immunotherapy has transformed the clinical outlook for many tumor subtypes its impact on PDAC has been disappointing to date. One factor in this regard may be the characteristic nature of the PDAC microenvironment which is associated with an intense desmoplastic reaction characterized by an abundance of cancer associated fibroblasts (CAF) (2). This may act to limit the access of immune effector cells to tumor (3,4), promote the infiltration of immune suppressive leucocyte populations and directly support tumor growth (5-8). The potential targeting of cancer associated fibroblasts in pancreatic cancer is now of considerable interest. However, studies from murine models have also shown that fibroblasts may play a protective role in limiting metastasis of the primary tumor and that targeting of stromal cells can increase tumor metastasis (9). As such, effective fibroblast-targeted therapies will need to be directed towards those subpopulations that are critical to support tumor growth. CAF display marked plasticity and can differentiate from a range of additional cell types under the influence of tumor-secreted factors (10) and their core transcriptional features can be difficult to define although it has been suggested that CAF populations with the features of myofibroblasts localize near to tumor within the PDAC microenvironment whilst inflammatory populations are observed more distal from tumor (11). Immunotherapy trials in PDAC indicate that multimodal approaches may be required for effective therapy. Therapeutic targeting of CAF populations is likely to require targeting of subsets in apposition to tumor cells that both support their growth and act to limit the tumor-specific immune response. Here we combine spatial transcriptomics with scRNA-Seq to define the transcriptome of tumor-proximal and tumor-distal fibroblasts within the PDAC microenvironment. Furthermore, these studies were correlated with clinical outcome and reveal that elevated podoplanin and HIF-1α expression were markers of poor outcome whilst expression of immunoregulatory genes correlates with favorable long-term response. These findings provide insight into stromal architecture in PDAC and could help to guide therapeutic approaches to target pro-tumorigenic fibroblast subsets.

## Results

### Spatially-defined stromal and immune regions can be characterized within the PDAC microenvironment

Histological slide sections were obtained from tumor biopsies of 24 patients with pancreatic ductal adenocarcinoma (PDAC) who had undergone surgical resection for localized disease. Thirteen patients had died of PDAC within 12 months of diagnosis (subsequently referred to as ‘poor response’) whilst 11 had survived for at least 36 months (‘good response’). Four-plex immunofluorescence staining was used initially to define major anatomical subregions of the tumor. Antibodies against pan-cytokeratin, α-SMA and CD45 identified epithelial, fibroblast and immune populations respectively whilst DAPI staining defined nuclear architecture. Regions in which stromal cells were adjacent to tumor (‘tumor-proximal stroma’), distant from tumor (‘tumor-distant stroma’) or enriched for CD45+ immune cells (‘immune enriched’) were then selected and 4 areas (‘regions of interest’; ROI) within each of these 3 domains were selected from each patient for assessment using the NanoString GeoMx Digital Spatial Profiler (DSP) platform (**Fig. 1**).

**Fig. 1.**
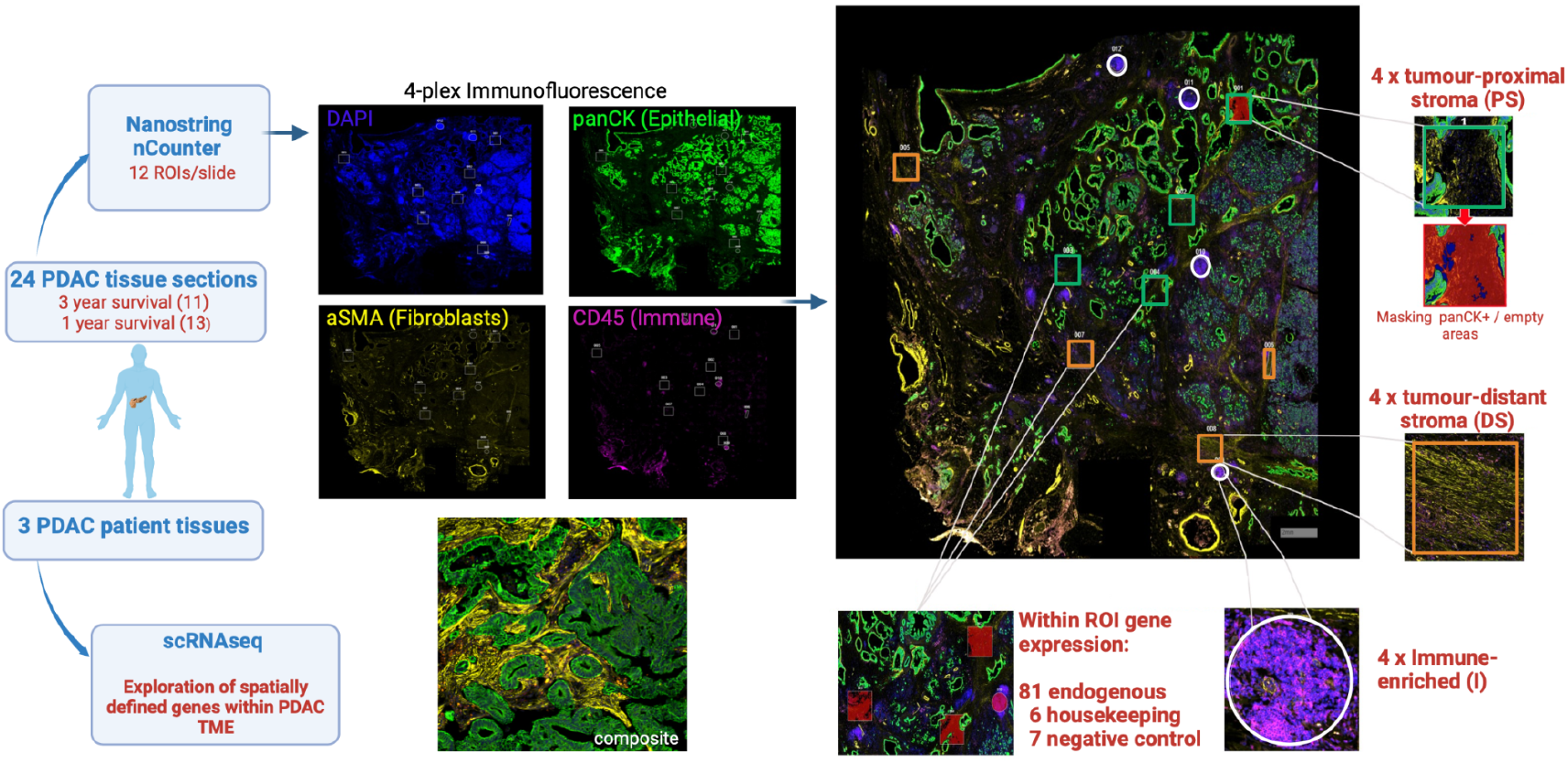
Schematic representation of experimental approach. Histological slides of 24 surgical resection specimens from patients with PDAC were stained with DAPI (‘nuclear’), anti-pan-CK (‘epithelial’), anti-α-SMA (‘fibroblast’) and anti-CD45 (‘immune’) to identify tumor cells and three other domains: ‘tumor-proximal stroma’ (PS), ‘tumor-distant stroma’ (DS) and ‘immune-enriched’ (I). NanoString’s Immuno-oncology RNA probe set plus a fibroblast-targeted custom panel of 10 RNA probes was used to interrogate the four areas (Regions of Interest; ROI) from each of the 3 domains on each slide using the NanoString GeoMx Digital Spatial Profiler (DSP) platform. The transcriptional profile of spatially-defined fibroblast proximal or distant cells was also extended using scRNA-Seq data from three additional PDAC samples.

94 RNA hybridization probes **(Supp. Table 1, Supp. Fig. S1 and S2)** for 81 endogenous, 6 housekeeping and 7 negative control genes were then applied to the 12 ROI from each patient, thus generating 288 transcriptional datasets. 6 pan-cytokeratin positive ROI were also selected to define the transcriptional profile of tumor cells. Hierarchical clustering of transcriptional datasets delineated immune and stromal regions with two immune profiles clustering separately due to differential expression of activatory and inhibitory immune genes. Cell-type expression profiles were consistent with immune or stromal origin (**Fig. 2A**). UMAP analysis broadly separated tumor, proximal stroma, distal stroma, and immune regions (**Fig. 2B**) although overlay of clinical outcome data did not reveal significant clustering.

**Fig. 2.**
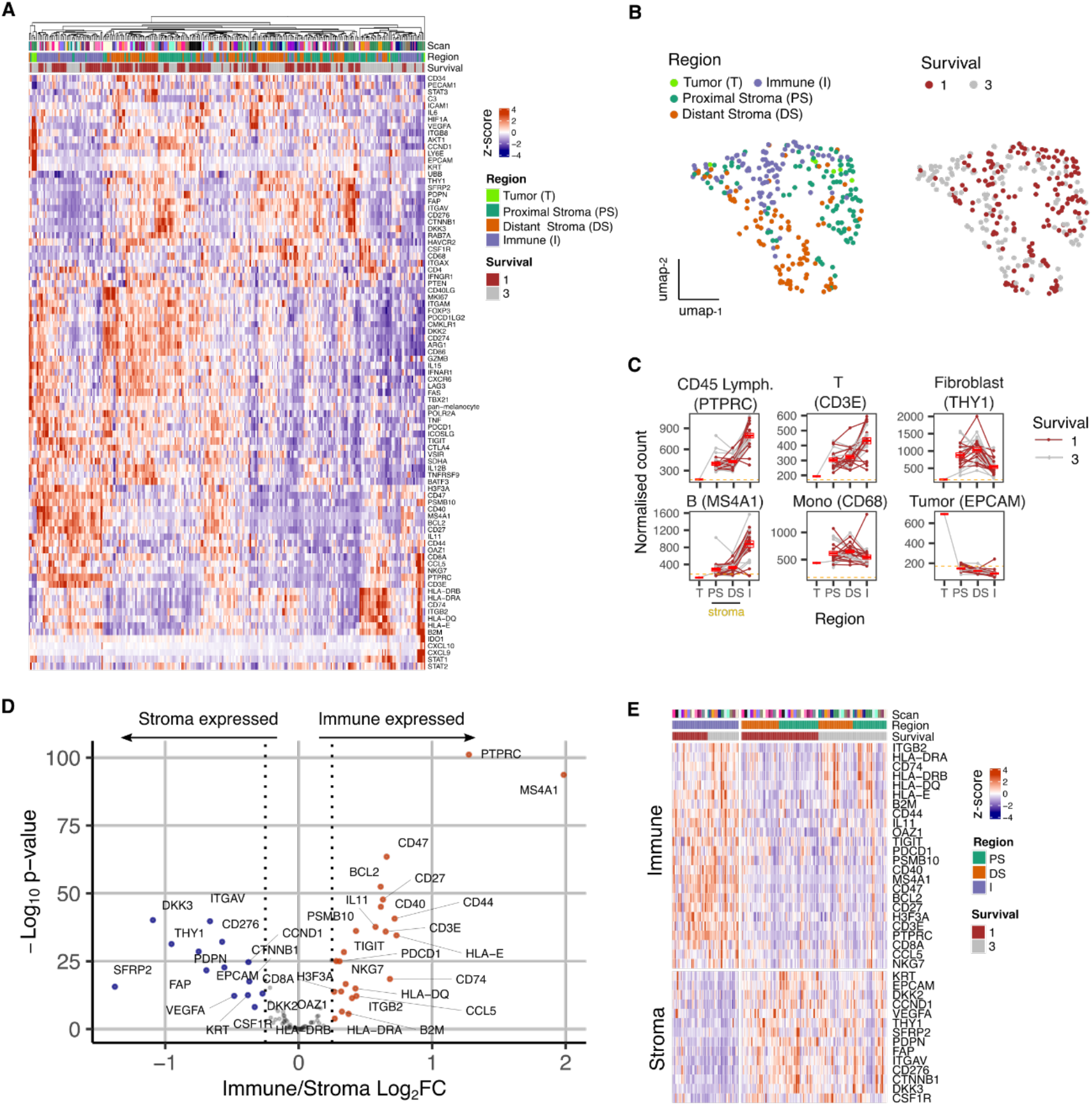
Overview of gene expression data from spatially-defined stromal and immune regions within the PDAC tumor microenvironment using NanoString GeoMx DSP. **A**. Expression profile of all endogenous probes across regions of interest (ROI) with hierarchical clustering of ROIs. **B**. UMAP embedding from normalized count data showing all ROIs overlaid with ROI-specific annotations of Region (Immune/Stroma/Tumor type) and Survival (1yr/3yr). **C**. Mean normalized count by region type for canonical high level cell type marker genes. Lines indicate regions from the same patient; dashed line represents mean background threshold from negative probes; Mean +/- SE of mean shown in red. **D**. Differential expression analysis identifies genes expressed differentially between Immune and Stroma ROIs. Colored points indicate differentially expressed genes (DEG) (BH adjusted p <0.05 & absolute log_2_FC > 0.25). **E**. Immune and Stroma expression signatures from DEGs identified in D.

Expression of cell lineage marker genes was then used to determine the relative localization of cell subsets within proximal-stromal, distal-stromal, immune or tumor regions of interest **(Fig. 2C)**. These confirmed localization of epithelial, fibroblast and lymphoid cells within the tumor, stromal and immune regions respectively whilst monocyte representation was equivalent within stromal and immune regions, consistent with broad infiltration within PDAC microenvironment. CD3E and MS4A1 expression indicated that stromal regions also contained smaller populations of infiltrating T and B cells (**Fig. 2C**). Distinct modules of genes co-expressed with lineage markers could also be identified and were consistent with cell type **(Supp. Fig. S3)**. Stromal regions expressed canonical fibroblast markers such as THY1, PDPN, FAP and THY1 whilst immune-specific genes such as PTPRC, CD3E and MS4A1 were present within Immune regions (**Fig. 2 D,E, Supp Fig. S4**).

To align the regional transcriptional landscape to specific cell subsets, RNA expression profiles defined by NanoString DSP analysis were mapped onto an additional scRNA-Seq dataset (**Supp. Fig. S5**) (Pearce et al. 2022; *in Press*). This revealed that, whilst the great majority of stromal-associated genes were expressed from fibroblasts, the expression of *CSF1R* within stroma was largely derived from myeloid cells, *CTNNB1* localized to endothelial cells and expression of *KRT* was identified as *KRT18* within epithelial cells (**Supp. Fig. S5C**).

### The transcriptional profile of stromal regions is strongly determined by proximity to tumor

We next went on to assess gene expression within stroma in relation to proximity to tumor (**Fig. 3**). Transcriptional profiles were seen to vary markedly between tumor-proximal or tumor-distal ROI. In particular, expression of *DKK3* and *PDPN* was markedly increased in stroma-proximal regions **(Fig. 3A,B,C)** and both are established markers of cancer-associated fibroblasts implicated in support of tumor growth (12-14). In contrast, *C3, SFRP2, STAT3, IL-6* and *THY1* expression was increased in tumor-distal stroma. C3 and SFRP2 were particularly elevated (**Fig. 3C**) and C3 expression was confirmed as fibroblast-derived by significant positive correlation with canonical fibroblast marker genes **(Supp Fig. S6)**. This is noteworthy given emerging importance for intracellular complement expression and the action of SFRP2 as a Wnt inhibitor. STAT3 and IL-6 expression is explained by their presence within inflammatory CAF whilst THY1 is commonly expressed on stem-like populations of fibroblasts (15). The stem cell marker CD34 was also expressed in this region (**Fig. 3A,B,C**). Immunohistochemical staining confirmed extreme polarization of podoplanin, DKK3 and C3 expression in relation to tumor proximity. Podoplanin was expressed on stroma that encased tumor whilst DKK3 expression was present both within tumor and tumor-proximal stroma. In contrast, expression of C3 was localized to distal stroma regions (**Fig. 3D**).

**Fig. 3.**
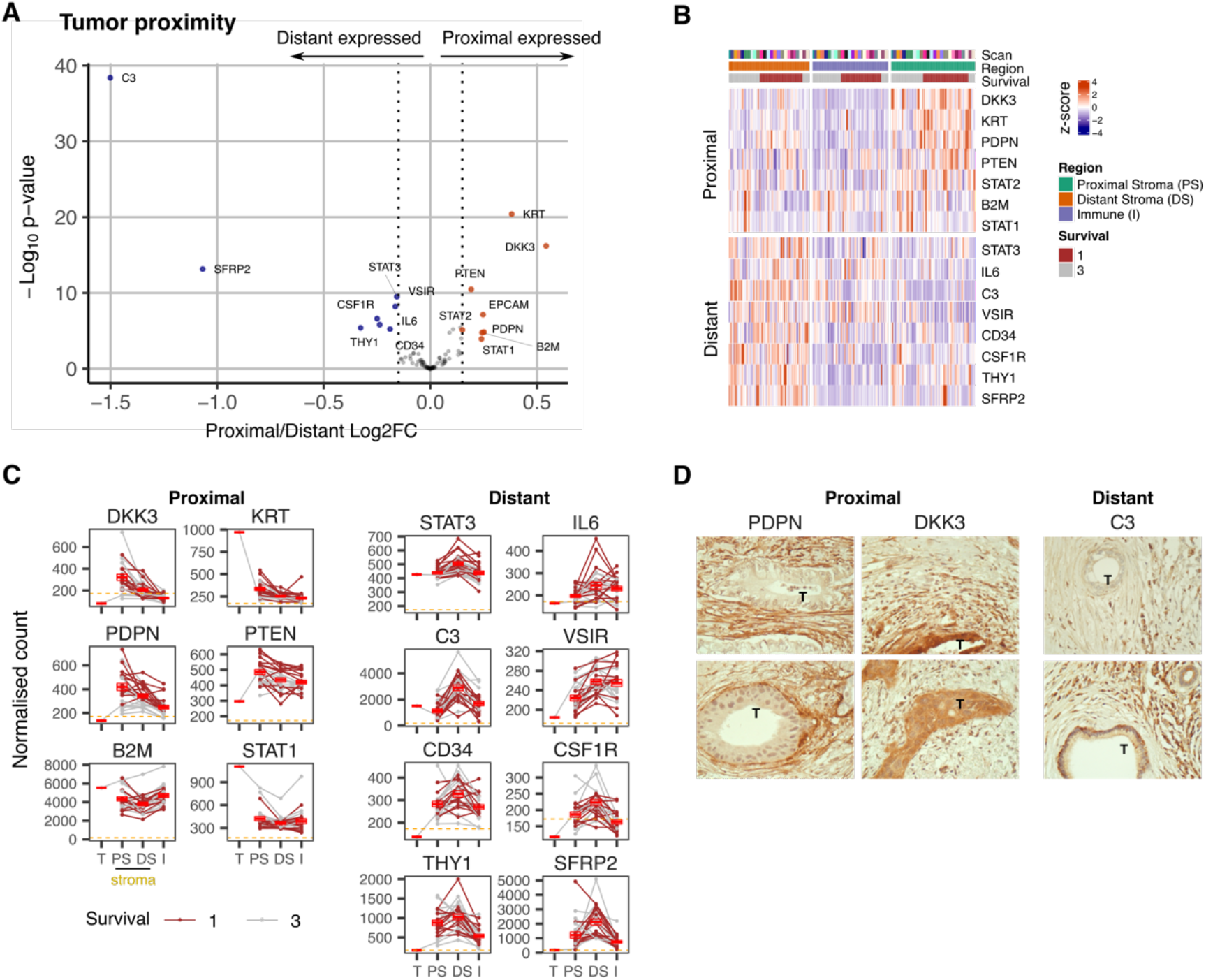
Expression signature of PDAC tumor-proximal and tumor-distal stromal cells. **A**. Differential expression of genes (DEG) within stroma regions proximal (P) or distal (D) from tumor. Colored points indicate differentially expressed genes (BH adjusted p <0.05 & absolute log_2_FC >0.25). **B**. Stroma proximity-to-tumor expression signature from DEGs identified in A. **C**. Relative expression of genes within four PDAC regions: Tumor (T), proximal-tumor stroma (PS), distal-tumor stromal (DS) and immune (I). Lines indicate paired regions from the same patient; dashed line represents mean background threshold from negative probes; Mean +/- SE of mean shown in red. Shown as within patient mean normalized count vs region type for DEG identified in A. **D**. Representative immunohistochemical staining of podoplanin, DKK3 and C3 proteins in relation to tumor cells (T) in PDAC tissue.

### Mapping of spatial transcriptional profiles on to scRNA-Seq reveals key biochemical pathways associated with proximal and distant fibroblasts

Given the profound influence of tumor apposition on the NanoString profile of fibroblasts we were interested to explore global fibroblast transcriptome in relation to spatial localization. The minimal NanoString gene set defining proximal and distant fibroblast subsets was therefore applied to the scRNA-Seq dataset (**Fig. 4A**) and two fibroblast clusters were observed from scRNA-Seq analysis and defined as sc-proximal and sc-distal populations due to their distinct proximal and distant gene expression signatures **(Fig. 4B,C)**. Transcriptional profiles were highly divergent between proximal compared to distal clusters with 47 genes differentially upregulated in the distal cluster and 36 genes differentially upregulated in the proximal cluster (**Fig. 4D**). sc-proximal clusters showed high expression of myofibroblast (myCAF) marker genes including *MMP11* and *HOPX* whilst the distal population was enriched for expression of genes associated with inflammatory CAF (iCAF) such as *CXCL12* and *CFD* **(Fig. 4E)**.

**Fig. 4.**
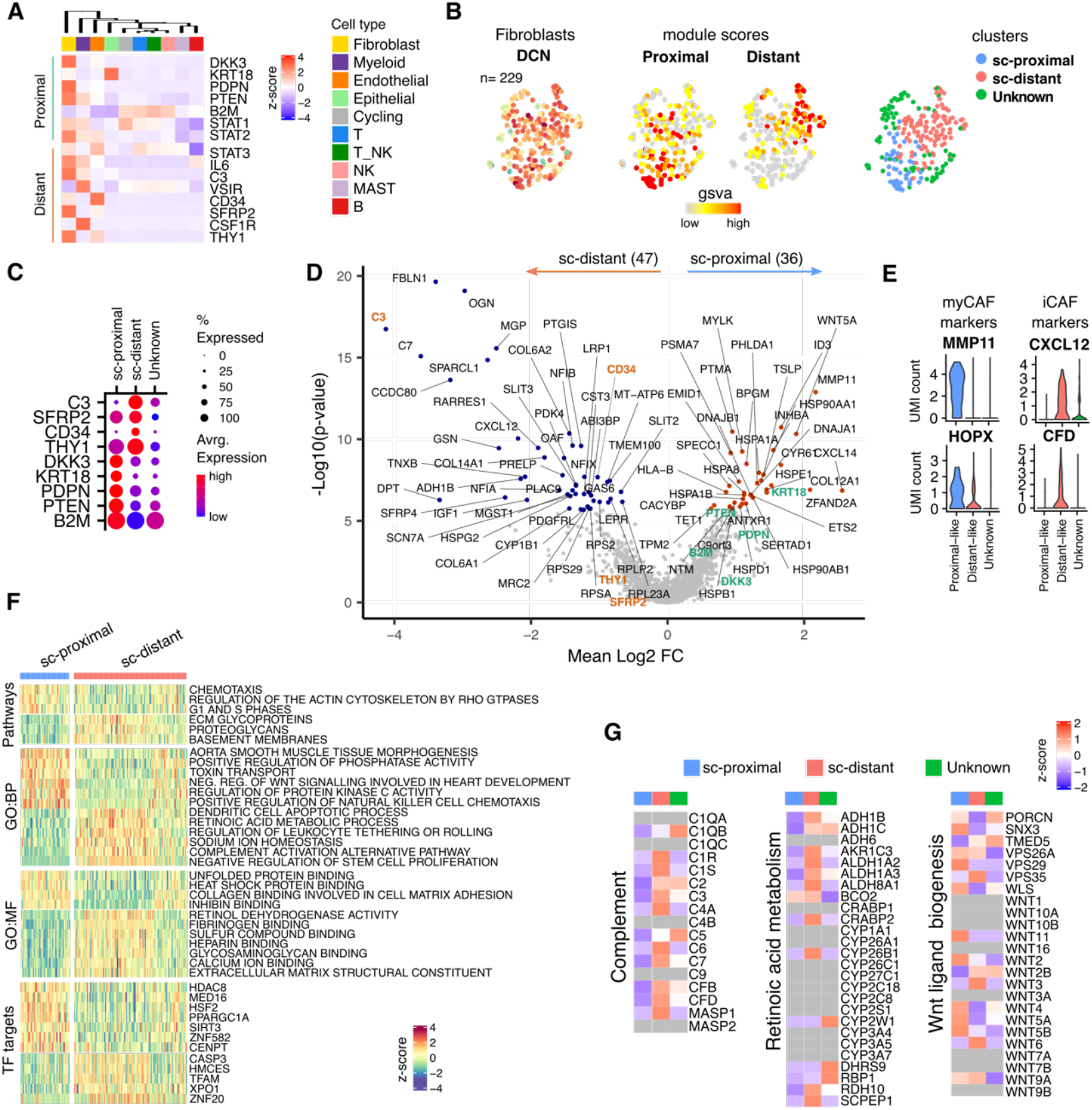
Proximal and Distal Fibroblast populations identified in PDAC single cell transcriptome data. **A**. Average expression of spatially defined tumor-proximal or tumor-distant stromal genes within cell types defined by scRNA-seq (n=3). **B**. UMAP embedding of scRNA-Seq data from fibroblasts overlaid with expression of the canonical fibroblast marker gene *DCN* and then the gene set variation analysis (GSVA) signature score for tumour proximal (*DKK3, PDPN, PTENSTAT2, B2* and *STAT1*) or tumour-distal (*STAT3, IL6, C3, VSIR, CD34, CSF1R, THY1, SFRP2)* associated stromal genes. Clustering based on unsupervised Louvain assignment. n=229 cells. **C**. Average cluster-wise expression profile of selected proximal and distant stroma associated genes as identified by spatial profiling. **D**. Differential expression analysis between sc-proximal and sc-distant fibroblast cells. Colored points indicate differentially expressed genes (BH adjusted p < 0.05 & absolute log_2_FC > 0.5). **E**. Violin plots depicting cluster-wise expression distribution of canonical myCAf and iCAF marker genes. **F**. GSVA score profiles identified as differentially enriched (BH adjusted p < 0.001) in sc-distant vs sc-proximal cells. **G**. Average within-cluster expression profile of Complement, Retinoic acid metabolism and Wnt ligand biogenesis gene sets. Grey = no detectable expression.

Differential pathway activity analysis showed that sc-proximal fibroblasts are enriched for cell division, chemotaxis and heat shock protein binding. Furthermore, they express a wide range of Wnt ligands including WNT5A, WNT11, WNT2, WNT5, WNT5A and WNT5B (**Fig. 4F,G**). In contrast, sc-distal fibroblasts mapped to pathways associated with generation of the extracellular matrix and negative regulation of stem cell proliferation (**Fig. 4F**). Also notable was expression of many members of the complement pathway as well as many genes associated with retinoic acid metabolism (**Fig. 4G**)

In order to assess the regulation of differential fibroblast transcription we next assessed relative expression of transcription factor targets in proximal and distal populations (**Fig. 4F**). This showed considerable divergence and reveals how spatially determined expression of transcription factors underpins differential fibroblast programming.

### Podoplanin and hypoxia predict poor outcome whilst high level expression of immune regulatory genes associates with superior clinical outcome

The study cohort had been selected to comprise patients with poor or good clinical outcome in order to identify potential spatial transcriptional correlates of disease progression. Poor outcome was defined as death within 1 year whilst patients with good outcome exhibited survival beyond 3 years **(Fig. 5)**.

**Fig. 5.**
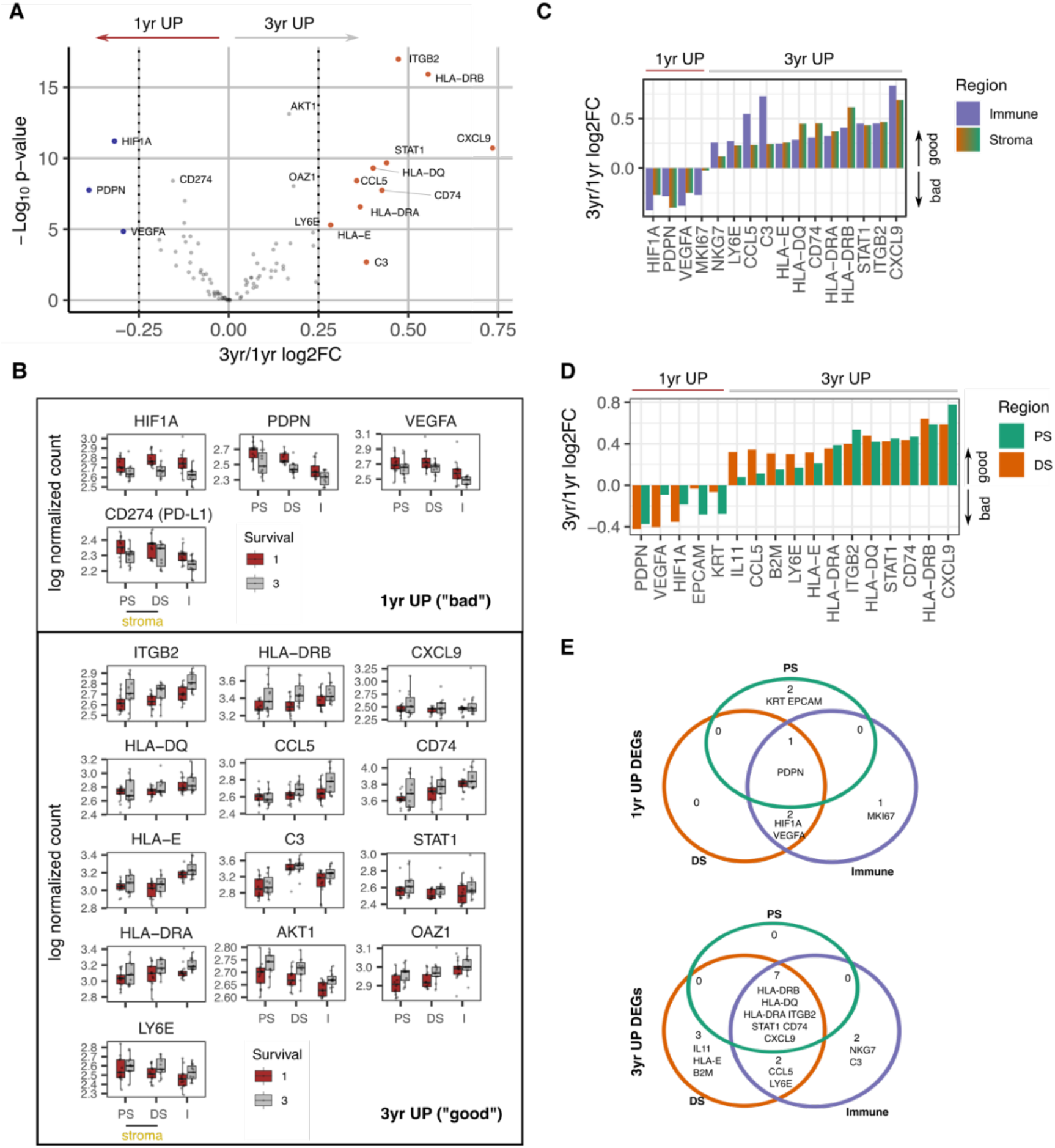
Survival expression signatures within spatially defined regions of PDAC. **A**. Differential gene expression from all regions in relation to poor (<1 year) or good (3+ year) survival. Coloured points indicate differentially expressed genes (BH adjusted p <0.05 & absolute log_2_FC > 0.25). **B**. Regional expression of survival-associated genes identified in A. Mean +/- SE of mean. PS=Proximal Stroma; DS=Distant Stroma; I=Immune. **C**. 3yr/1yr fold change in expression of survival-associated genes within Immune and Stroma regions. (BH adjusted p < 0.05 absolute log_2_FC > 0.25). **D**. 3yr/1yr fold change in expression of survival-associated genes within Tumor-Proximal and Tumor-Distal regions. (BH adjusted p < 0.05 & absolute log_2_FC > 0.25). **E**. Venn displaying overlaps of 3yr vs 1yr survival DEGs (BH adjusted p < 0.05 absolute log_2_FC > 0.25) within tumour-proximal stroma (PS), tumour-distal stroma (DS) and Immune (I) regions

Overall gene expression profiles were initially compared between these two groups in order to define prognostic transcriptional signatures. High level transcriptional expression of PDPN, HIF1A,PDL1 (CD274) and VEGFA were associated with poor clinical outcome (**Fig. 5A, Supp Fig. S7B**). Ten genes were upregulated in patients with survival beyond 3 years and were characterized predominantly by immune activation with increased expression of MHC class I and class II, complement C3 and chemokines CCL5 and CXCL9. The integrin ITGB2 (CD18) and STAT1 also showed increased expression in this group.

The spatial expression of these prognosis-associated genes was then assessed across the two risk groups **(Fig. 5B)** and the relative expression was also determined in immune and stromal regions **(Fig. 5C,D)**. Patients with a poor outcome expressed PDPN broadly across the tumor microenvironment whilst HIF-1α and VEGF expression also extended into the distal stromal and immune regions and likely indicates more extensive hypoxia in this subgroup. Good prognosis was associated with broad expression of most immunostimulatory and immunoregulatory genes whilst expression of IL-11 and HLA-E was focused within distal stroma **(Fig. 5C,D,E)**. Expression of complement C3 and NKG7, a regulator of cytotoxic granule release (16) within the immune region was also enhanced in patients with good clinical outcome..

## Discussion

Increased understanding of the pancreatic ductal adenocarcinoma microenvironment is essential for the development of targeted therapies. Here we combined spatial and single cell transcriptomic analysis to interrogate patterns of cellular transcription in relation to tumor proximity and related this to clinical outcome. This reveals spatially-determined transcriptional programming of fibroblasts with potential opportunities for therapeutic development.

Proximity to tumor was seen to be a strong determinant of transcriptional activity of stromal cells. In particular, DKK3 and PDPN were both increased markedly on tumor-proximal cells. PDPN expression is strongly enhanced on cancer-associated fibroblasts in PDAC (14) and high expression levels are correlated with poor prognosis. DKK3 is a Wnt regulator and is emerging as a potentially important therapeutic target (12). A range of genes showed increased expression within stromal populations distal from tumor including the C3 component of complement and SFRP2, a soluble modulator of Wnt signaling. CD34, a marker of stromal stem cells, was also expressed more highly in this region and may indicate spatial differentiation of stromal cells towards the tumor.

Integration of spatially-defined transcription signatures with single cell RNA-Seq data allowed development of a transcriptional atlas of proximal and distal stromal cells. Tumor-proximal populations displayed features typical of myofibroblasts whilst more distal populations had an inflammatory profile, in line with previous reports (11). Myofibroblast markers included the potent pro-tumorigenic chemokine CXCL14 (17) and WNT5A which may contribute to the differentiation of adipocytes to CAFs (18). Indeed, Wnt ligand signaling plays a key role in PDAC progression and therapeutic resistance and tumor-proximal fibroblasts are seen to be strong contributors to the Wnt ligand pool with high level expression of the ligands WNT5A, WNT11, WNT2, WNT5, WNT5A and WNT5B. Transcriptional regulation of cell division was also increased suggesting enhanced proliferation of stromal cells when locally exposed to tumor. In contrast, stromal cells located more distally from tumor retained functions such as generation of extracellular matrix protein. Striking expression of a wide range of complement proteins was also seen at this site. CAF populations expressing complement proteins have been observed previously in PDAC (19) and overlap with the transcriptional profile of inflammatory CAF. The physiological role of intracellular complement expression is receiving considerable interest with evidence that it may impact on immune surveillance in pre-clinical models (20). A further finding of note was increased expression of a range of genes associated with retinoids and is noteworthy given that vitamin A-containing lipid droplets are enriched within quiescent pancreatic stellate cells in close proximity to the basal aspect of pancreatic acinar cells (21). Indeed, patients with PDAC are often vitamin A deficient whilst retinoic acid treatment can suppress stellate cell proliferation with associated reduction in Wnt-β-catenin signaling and localized tumor apoptosis (22). ATRA treatment has been shown to be tolerable in patients with advanced disease and is under investigation in phase I trials (23). Distal populations were also enriched for expression of genes associated with negative regulation of stem cell proliferation and may indicate a potential role for cells within this environment in limiting tumor cell progression.

CAF populations exhibit extreme plasticity and factors such as IL-1 and TGF-β are emerging as important mediators of local phenotype (24). Analysis of relative transcription factor binding expression within tumor-proximal or distal stroma identified substantial differences in transcription factor activity at the two sites. A wide range of transcription factors were differentially expressed and indicate the importance of local cellular environments in exploiting the transcriptional plasticity of fibroblasts.

The study cohort had been selected to include patients with poor or good clinical outcome, based on survival below 1 year or above 3 years, respectively. High level expression of podoplanin, HIF-1α and VEGF were associated poor outcome. The negative prognostic impact of podoplanin expression in PDAC has been documented previously and podoplanin-positive stromal cells enhance invasion and proliferation of tumor cells. However, downregulation of podoplanin expression does not reverse this effect indicating an important role for additional pathways within this population (14). Expression of HIF-1α is reflective of the hypoxic environment within PDAC tumors and indicates that the intensity of hypoxia is an independent determinant of clinical outcome (25-27). Indeed, spatial extension of HIF-1α expression into distal stroma and immune microenvironments was an additional risk factor and indicates that the breadth of hypoxia is of prognostic importance. HIF-1α expression in PDAC is associated with a range of features including enrichment of glycolysis, modulation of mTORC1 and MYC signaling, and immune suppression (28,29). As such, this represents a challenging tumor subgroup for therapeutic intervention although the introduction of HIF-1α inhibitors offers encouragement in this regard (30). Hypoxia is also likely to explain increased levels of VEGF expression in patients with poor prognosis. VEGF-targeted therapies have not shown significant utility in PDAC but could potentially be considered as part of a multi modal therapeutic approach (31).

In contrast, many immunoregulatory and immunostimulatory genes were increased in patients with good prognosis and concur with studies showing that the extent of lymphocytic infiltration is a favorable indicator for outcome. Liudahl et al recently used chromogen-based multiplexed immunohistochemistry (mIHC) to generate an atlas of leucocyte contexture within PDAC (32) and extended prognostic utility to immune subpopulations. It was noteworthy that elevated expression of HLA class II genes was seen in patients with longer term survival and as this association extended into stromal regions it may indicate an important role for HLA-DR+ antigen-presenting CAF (ApCAF) populations (33). Expression of complement protein C3 was associated with good clinical outcome and indicates that this pathway can also help to contain tumor growth (34) despite early indications of a potential pro-tumorigenic role (20). Indeed, the beneficial effect of ApCAF in lung cancer is mediated partially through expression of complement proteins which rescue intratumoral T cells from exhaustion (35) and this may provide a unifying explanation for the prognostic value of HLA class II and complement expression in this study.

Expression of IL-11 within distal stroma was also a positive prognostic sign and is noteworthy given a previous report of a similar association with elevated serum concentrations (36). IL-11 is an inflammatory protein within the IL-6 family and as such further analysis of the mechanisms by which it can help to contain PDAC development would be valuable. High level expression of NKG7 within immune regions was also beneficial and, given its central role in regulation of cytotoxic granule release (16), this is noteworthy given its emerging role as a predictive factor in response to checkpoint protein inhibition (37). Overall, the transcriptional correlates of good prognosis clearly identify immunological processes as the central determinant of clinical outcome and augur well for therapeutic interventions that can unmask this immune potential. Immune checkpoint inhibition has been largely unsuccessful for this patient subgroup but there is clearly latent immunogenicity within the PDAC microenvironment and the use of agonistic anti-CD40 antibodies has shown promise in clinical studies (38).

In conclusion, we find that transcriptional activity of stromal subsets is strongly regulated by their relative proximity to tumor and define the transcriptional landscape in relation to spatial localization. Hypoxia is a correlate of poor outcome whilst approaches to enhance the inflammatory environment of distal stroma could offer approaches to improve the clinical outcome for this patient group. Indeed, successful therapy for PDAC may require multi-modal approaches and recent pre-clinical evidence of a synergistic activity of HIF inhibition with immune checkpoint blockade in pre-clinical cancer models is one such area of interest (39).

## Materials and Methods

### Participants

FFPE tissue from 25 treatment-naïve patients undergoing pylorus-preserving pancreaticoduodenectomy (PPPD) who presented with localized disease were selected for this study. The study design was approved by ethical approval by Birmingham Local Research Ethics Committee (REC 16/WM/0214) and the HBRC Biobank ethics 18-304. Written informed consent was obtained from patients, and studies were conducted in accordance with the Declaration of Helsinki.

### Sample processing

FFPE tissue blocks were sectioned at 5-μm thickness, deparaffinized and rehydrated using conventional methods. The slides were profiled using NanoString GeoMx Digital Spatial RNA Profiling (DSP) platform through the Technology Access Program (TAP) by Nanostring (Seattle, WA, USA). Briefly, immunofluorescent antibody staining was performed with tissue morphology markers (Pan-CK, CD45, α-SMA and DAPI). In parallel, slides were stained with a panel of photocleavable RNA probes. Custom regions of interest (ROI) were selected based on these markers to generate specific domains including ‘tumor-proximal stroma’, tumor-distant stroma’ and ‘immune enriched’ areas. Four ROI were selected for each domain per slide. UV-cleavable probes within each ROI were liberated by UV light, hybridized to optical fluorescent barcodes then counted on the Ncounter to determine the absolute number of mRNA transcripts.

### NanoString nCounter data analysis

Raw NanoString nCounter data expression matrix was processed following the normalization and quality control procedures as described elsewhere (40). Due to a redundancy in the tags used for both IFNG and ACTA2, data from these probes had to be removed from further analysis. Correlations of housekeeping gene expression across all ROIs were assessed to select the most correlated housekeeping probes H3F3A and UBB to use for downstream normalization **(Supp. Fig. S2A**). Unwanted variation was removed using the R package RUVSeq (41). Firstly, distributional differences were scaled between lanes using upper-quartile normalization then unwanted technical factors were estimated in the resulting gene expression data with the RUVg function selecting H3F3A and UBB as the negative control genes and the number of dimensions of unwanted variation to remove set to 1. A variance stabilizing transformation of the original count data was computed using DESeq2 (42) and estimated unwanted variation was removed using the removeBatchEffects function from limma (43). RLE plots were used to detect any potential outliers before and after normalization (**Supp. Fig. S2B**).

Differential expression analysis was conducted to compare Immune vs stromal regions, 3 yr vs 1 yr survival and tumor-proximal vs tumor-distant stromal regions using DESeq2, adjusting for multiple testing with Benjamini-Hochberg (BH) procedure. Differentially expressed genes were determined by BH adjusted p < 0.05 and absolute log2FC > 0.25.

Dimensionality reduction by Uniform Manifold Approximation and Projection (UMAP) was performed on the normalized counts matrix with the umap R package and ggplot2 utilized for plotting. Heatmap visualizations were generated using the ComplexHeatmap package. Pearson correlation was calculated and plots generated using ggpairs and ggcorr functions from the R package GGally.

### scRNA-Seq data analysis

Genes of interest identified from nCounter data analysis were further explored for their expression profiles in single cell RNA sequencing data of cells within the tumor microenvironment of 3 PDAC patients (REF Pearce et al., 2023; in Press).

#### Raw read data processing

Raw reads were processed using CellRanger (10X Genomics, v3) functions mkfastq and count. Raw bcl files were converted to fastq and aligned to the human reference genome GRCh38. Gene expression matrices for each patient were analyzed by R software (v3.6). Data pre-processing, QC, dimensionality reduction, clustering and subsequent downstream analysis was performed using the Seurat package (v3.1.1).

#### Data integration and clustering

Data from 3 PDAC patient samples was integrated following Seurat SCTransform Integrate Data workflow using the top 3000 most variable genes as integration features. Principal Component Analysis (PCA) was applied and Uniform Manifold Approximation and Projection (UMAP) embedding determined using PCs 1:20. For unsupervised clustering, a shared nearest neighbour graph based on Euclidean distance in PCA space was constructed using Seurat FindNeighbours function and the modules within this graph representing clusters were identified using the 17ouvain algorithm with Seurat FindClusters.

To annotate clusters with high-level cell type, canonical cell type marker gene expression level was assessed.

#### scRNA-Seq Fibroblast data analysis

Transcriptome data was subset taking Fibroblast cells only and unsupervised clustering re-applied on Fibroblasts alone. Expression profile of stromal expressed genes identified from the nCounter dataset to be associated with tumor proximal or tumor-distant regions was assessed within the Fibroblast scRNA-Seq data. These tumor-proximal and tumor-distant gene signatures were scored using GSVA to assess likely tumor-proximal and tumor-distant fibroblasts. Expression profiling and GSVA signature scoring were used to annotate fibroblast subpopulations identified through clustering as “Proximal-like” and “Distant-like”.

To expand the pool of possible transcriptional markers for tumor proximal and tumor distant fibroblasts, differential expression analysis was conducted comparing Proximal-like and Distant-like clusters using findMarkers with MAST option (test.use=“MAST”), which uses a hurdle model tailored to scRNA-Seq data. MAST is a two-part GLM that simultaneously models how many cells express the gene by logistic regression and the expression level by Gaussian distribution (44). Differential expression testing was performed using the likelihood ratio test. Differentially expressed genes were determined by Benjamini Hochberg adjusted p < 0.05 and absolute log2FC > 0.5.

## Notes

**Funding** This research was supported by a Cancer Research UK (CRUK) Programme Grant CRUK-A21135.

**Competing Interests** The authors declare no potential conflicts of interest.

### Competing Interest Statement

The authors have declared no competing interest.

