## Supplementary Information for "Spatial determination and prognostic impact of the fibroblast transcriptome in pancreatic ductal adenocarcinoma"

**Supplementary Table 1. nanoString nCounter** **RNA hybridisation probeset**

| **Immunoncology core** | | | | **Fibroblast** | **Housekeeping** | **Negative** |
| --- | --- | --- | --- | --- | --- | --- |
| **L12B** | **IL6** | **CD11c** | **CXCR6** | **THY1** | **H3F3A** | **Neg Prb 1** |
| **IFNG** | **IFNGR1** | **CD11b** | **CTLA4** | **PDPN** | **SDHA** | **Neg Prb 2** |
| **STAT2** | **IFNAR1** | **4-1BB** | **CD40LG** | **CD34** | **POLR2A** | **Neg Prb 3** |
| **SDHA** | **ICAM1** | **TNF** | **PDL1** | **HLA-DRA** | **UBB** | **Neg Prb 4** |
| **PTEN** | **EPCAM** | **TIGIT** | **CD27** | **C3** | **RAB7A** | **Neg Prb 5** |
| **PECAM1** | **B2M** | **TBX21** | **BATF3** | **DKK3** |  | **Neg Prb 6** |
| **UBB** | **pan-Melanocyte** | **PDL2** | **VISTA** | **SFRP2** |  | **Neg Custom** |
| **OAZ1** | **Multi KRT** | **PD1** | **VEGFA** | **FAP** |  |  |
| **POLR2A** | **CXCL10** | **LAG3** | **STAT3** | **IL11** |  |  |
| **LY6E** | **CSF1R** | **IL15** | **STAT1** | **ACTA2** |  |  |
| **ITGB8** | **CD47** | **GZMB** | **CD45** | **OAZ1** |  |  |
| **ITGB2** | **CD44** | **FOXP3** | **PSMB10** |  |  |  |
| **ITGAV** | **CD40** | **DKK2** | **NKG7** |  |  |  |
| **CD20** | **CD8A** | **KI67** | **CD86** |  |  |  |
| **IDO1** | **CD68** | **ICOSLG** | **CD4** |  |  |  |
| **HLA-E** | **CD3E** | **HLA-DRB** | **B7-H3** |  |  |  |
| **HLA-DQA1** | **CCND1** | **HIF1A** | **CCL5** |  |  |  |
| **Tim3** | **BCL2** | **FAS** | **ARG1** |  |  |  |
| **CXCL9** | **AKT1** | **CTNNB1** | **RAB7A** |  |  |  |
| **CMKLR1** | **CD74** |  |  |  |  |  |

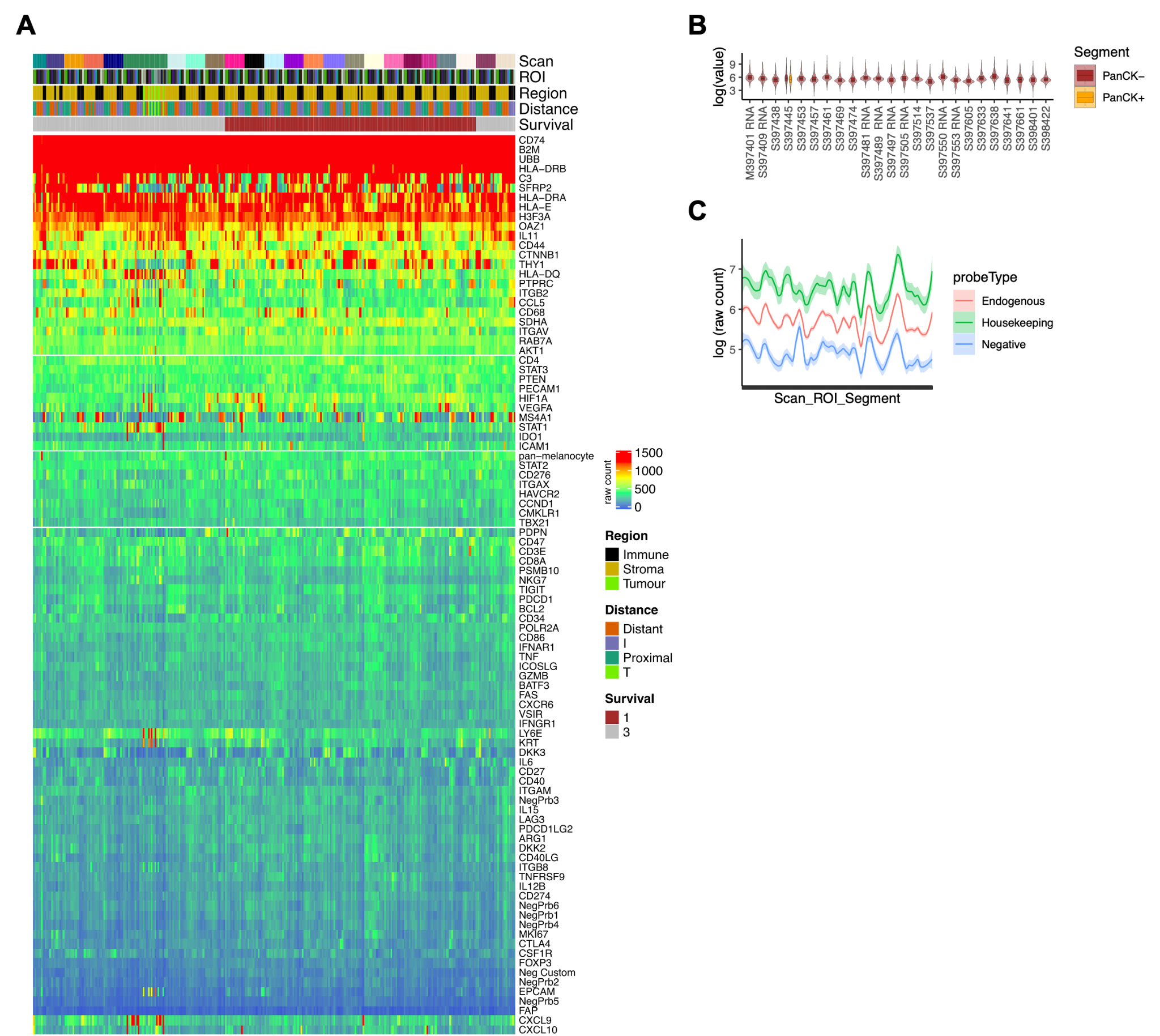

**Supplementary Figure S1. Raw count expression profiles.** A, Heatmap of raw expression profile for the complete probeset (colour scale= raw count). B, Per patient (scan) expression distributions for PanCK+ (tumor) and PanCK- (non-tumor) regions of interest. C, Per patient (scan) expression profiles (mean +/- SE) for Endogenous, Housekeeping and Negative control probesets.

**Supplementary Figure S2. Housekeeping gene correlations and data normalisation.** A, Pairwise correlation scatter plots for 6 housekeeping control probes. Raw housekeeping gene counts from all regions of interest (ROIs) were correlated against each other. Plots display the scatter, distribution histogram and Pearson correlation coefficient and significance (***p<0.001). B, Relative log expression plots of raw count data and post normalization with the top two most-correlated housekeeping genes (H3F3A and UBB) to remove unwanted variation (RUV method).

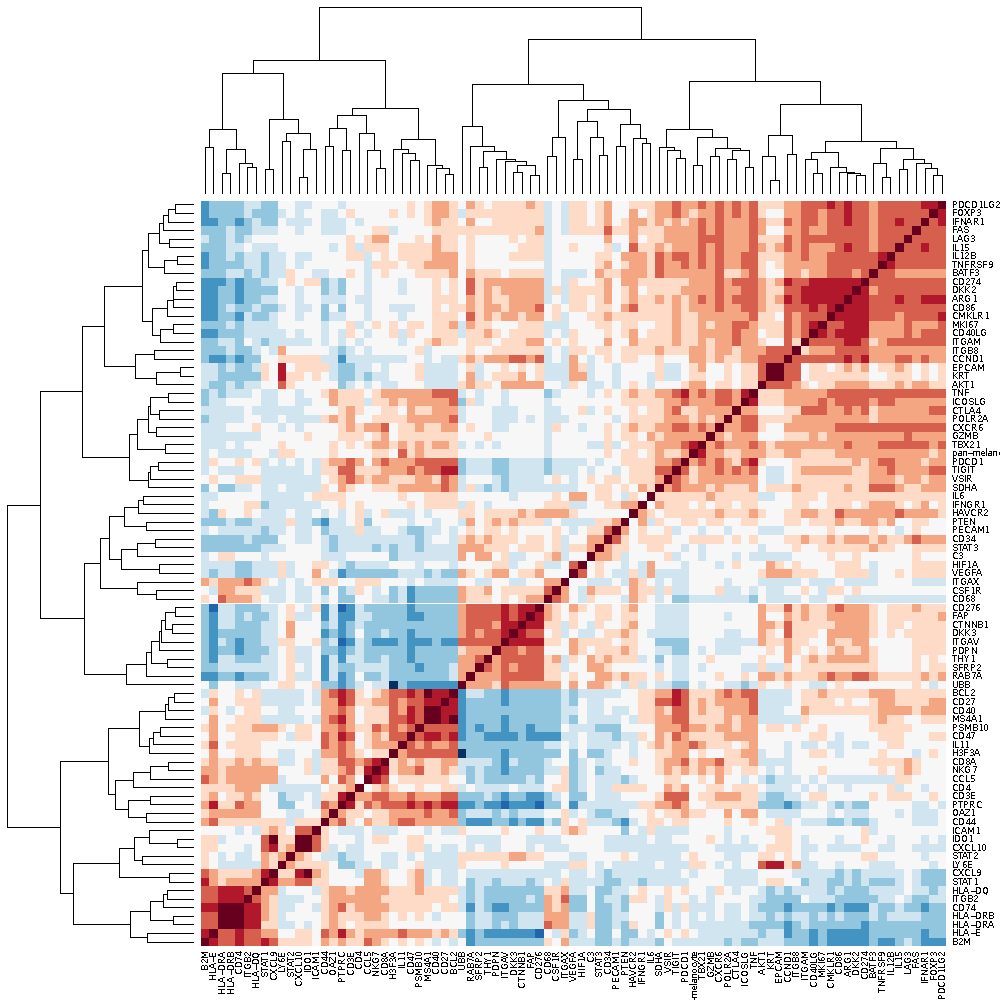

**Supplementary Figure S3. Correlation Matrix of endogenous probes.** Pairwise correlations of normalized gene count data represented as a matrix of Pearson correlation coefficients.

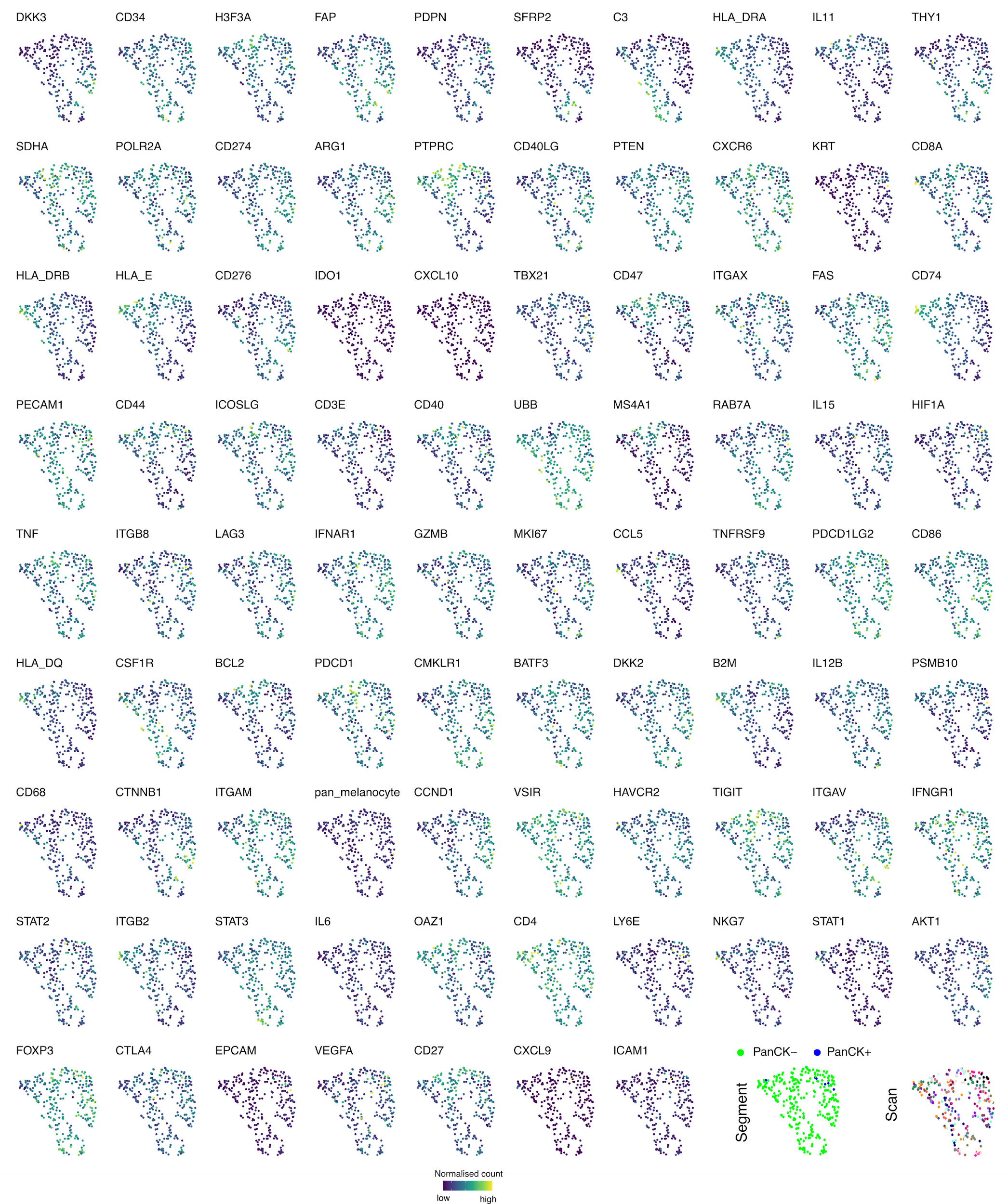

**Supplementary Figure S4. Individual gene expression profiles on UMAP embeddings of all regions of interest (ROIs).** UMAP embedding generated from normalized count data of all probes in all REIs overlaid with ROI-specific normalized gene count.

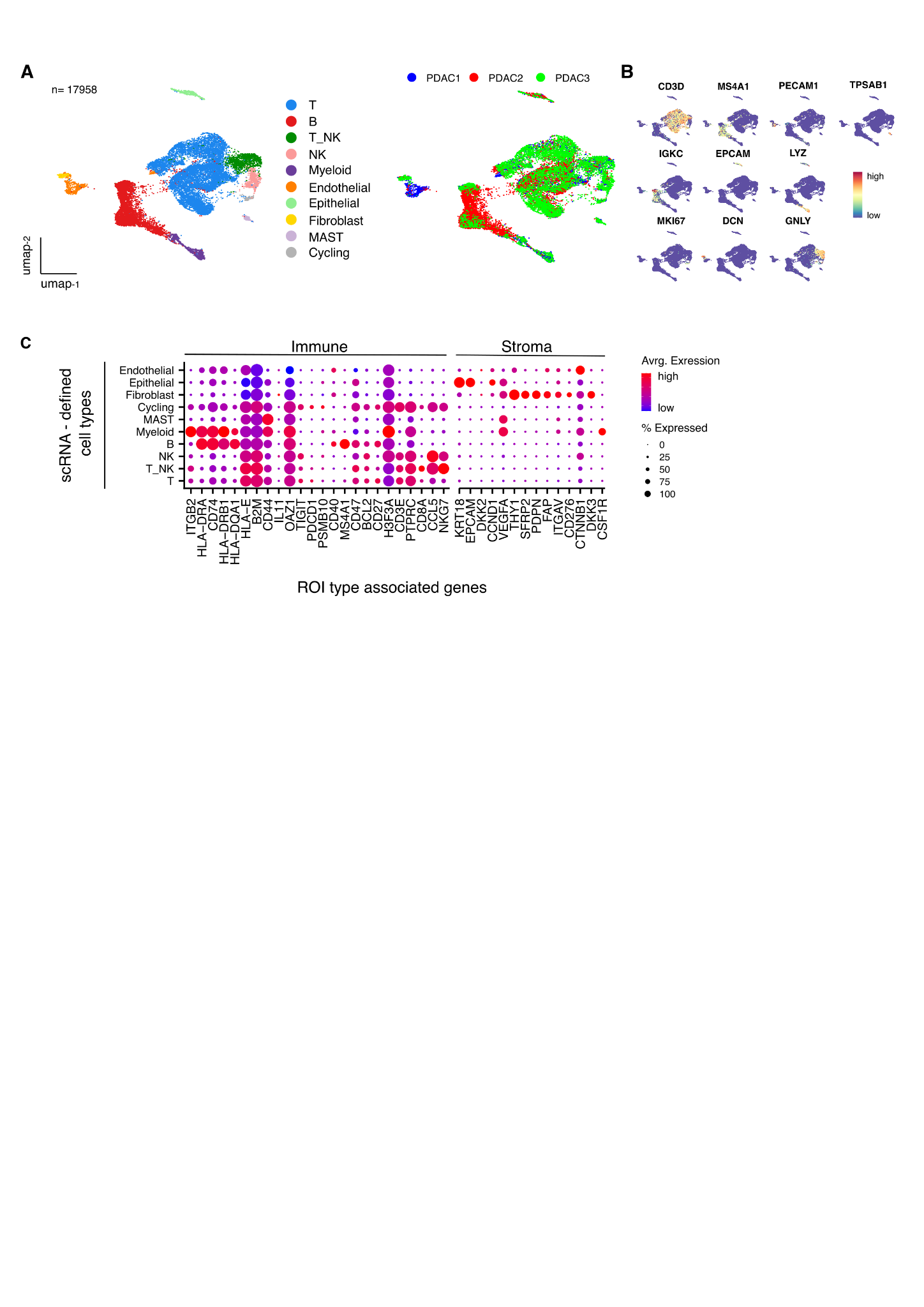

**Supplementary Figure. S5. High level cell type contexture of PDAC tumor microenvironment.** A (left), UMAP embedding of transcriptional data for 17,958 cells in the tumor microenvironment of 3 x PDAC patient tissue samples overlaid with high level cell type identified from unsupervised clustering. A (right), UMAP embedding overlaid with source sample identifier. B, UMAP embedding overlaid with the expression profile of canonical high level cell type marker genes. C. Average scRNA expression profile of spatial profiling defined Immune and Stroma ROI type-associated genes within high level cell types defined by scRNA-seq (n=3 PDAC samples).

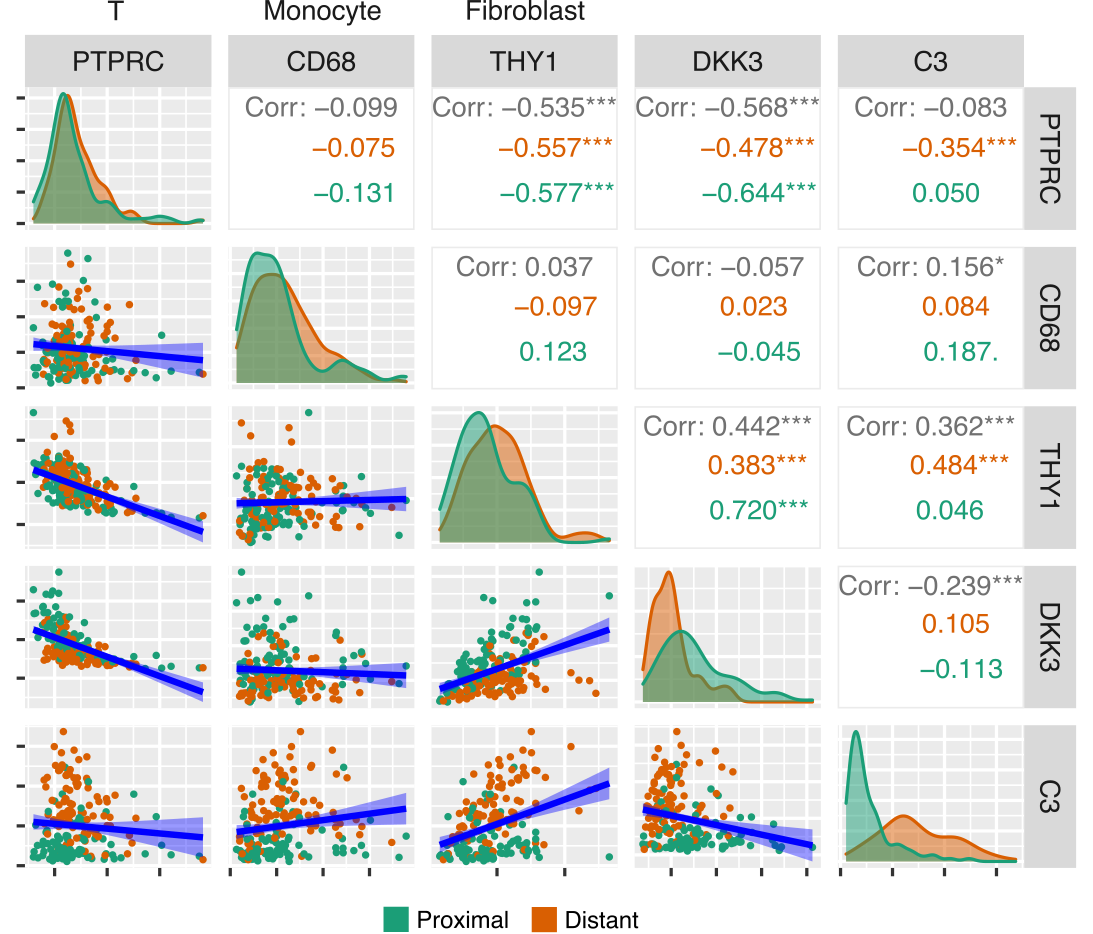

**Supplementary Figure. S6. Correlations of proximity specific markers DKK3 and C3.** Pairwise Pearson correlations of proximity specific markers C3 and DKK3 against marker probes for T, Monocyte and Fibroblast cells. Colour indicates data source; proximal or distant ROIs.

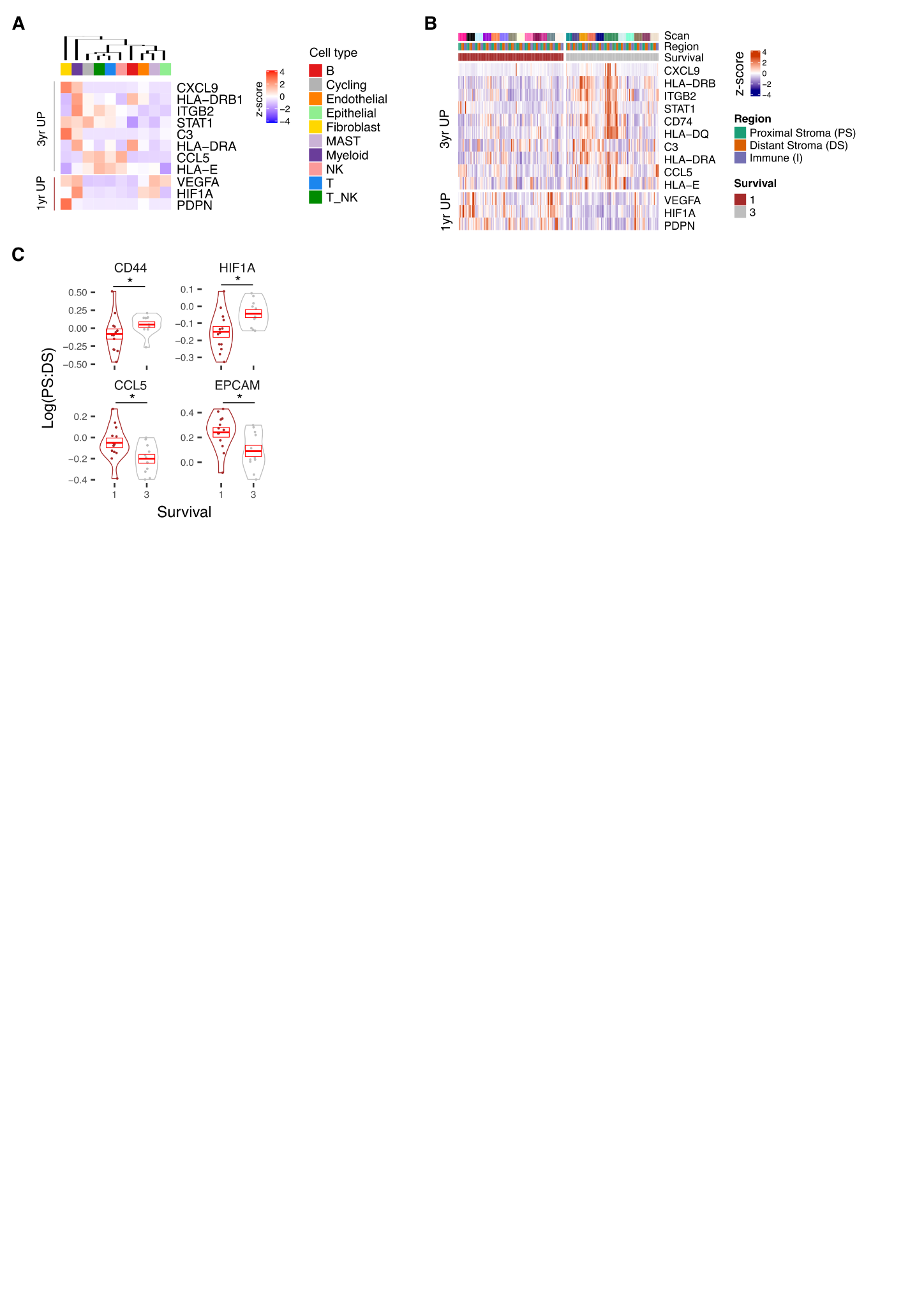

**Supplementary Figure S7. Profile of Survival expression signatures within PDAC TME.** A, Average scRNA expression profile of Survival associated genes within high level cell types of scRNA-seq dataset (n=3). B, Expression profile of Survival associated genes identified by 3yr vs 1yr differential expression analysis. C, Proximal Stroma (PS): Distant Stroma (DS) ratios stratified by survival outcome for selected genes with significant difference in ratio when stratified by survival.
